# Imputation of cancer proteomics data with a deep model that learns from many datasets

**DOI:** 10.1101/2024.08.26.609780

**Authors:** Lincoln Harris, William S. Noble

## Abstract

Missing values are a major challenge in the analysis of mass spectrometry proteomics data. Missing values hinder reproducibility, decrease statistical power for identifying differentially expressed (DE) proteins and make it challenging to analyze low-abundance proteins. We present Lupine, a deep learning-based method for imputing, or estimating, missing values in tandem mass tag (TMT) proteomics data. Lupine is, to our knowledge, the first imputation method that is designed to learn jointly from many datasets, and we provide evidence that this approach leads to more accurate predictions. We validated Lupine by applying it to TMT data from *>*1,000 cancer patient samples spanning ten cancer types from the Clinical Proteomics Tumor Atlas Consortium (CPTAC). Lupine outperforms the state of the art for TMT imputation, identifies more DE proteins than other methods, corrects for TMT batch effects, and learns a meaningful representation of proteins and patient samples. Lupine is implemented as an open source Python package.

## 1 Introduction

In spite of significant advances in instrument technology and sample preparation, missingness remains a challenge in quantitative mass spectrometry (MS) proteomics.^1, 2^ “Missingness” refers to peptides that are present in the analyte but for various technical reasons including co-elution, electrospray competition and inability to confidently assign peptide spectrum matches, do not have an associated quantitative value.^1, 2^ Missingness hinders reproducibility and reduces statistical power, making it difficult to compare across runs or experimental conditions. In addition, missingness can make it challenging to glean information from low-abundance peptides, which are important for disease etiology and progression in a number of contexts.^3, 4, 5^

Here we focus on the specific challenge of missingness in tandem mass tag (TMT) proteomics. TMT data are collected using data-dependent acquisition, in which missingness can mainly be attributed to the fact that precursors are stochastically selected for fragmentation and quantification, leading to peptides that are quantified in one run but not the next. Missingness is especially pronounced for large-scale, multi-batch TMT experiments, in which the number of missing values scales logarithmically with the number of TMT batches or “plexes”.^6^ Nevertheless, TMT offers unparalleled quantitative accuracy and has recently been used for large-scale studies of cancer and neurodegenerative disease.^7, 8, 9, 10, 11^ For example, TMT was used by the Clinical Proteomics Tumor Atlas Consortium (CPTAC) project^11, 8^ to analyze more than 1,000 clinical patient samples from ten types of cancer.

Empirically, missing values are not distributed entirely randomly but instead are typically correlated with the intensity of the peptide.^12^ In general, missing values may be either missing completely at random (MCAR) or missing not at random (MNAR).^13^ In the case of MCAR there is no relationship between underlying variables and the likelihood of an observation to be missing, whereas for MNAR there is. In MS proteomics, missing values tend to be MNAR because the likelihood that a peptide is missing depends on its intensity.^12^ Low-intensity peptides are more likely to be missing, although medium- or high-intensity peptides are occasionally missing as well.^12^

Imputation is an analytical solution to missingness. “Imputation” refers to using statistical or machine learning procedures to estimate missing values based on the observed values alone. Imputation is routinely used to handle missingness in data from microarrays,^14^ single-cell transcriptomics,^15, 16^ epidemiology,^17^ and astronomy.^18, 19^

Many methods exist for proteomics imputation; however, each of them has significant limitations. For example, many of these methods have been borrowed from microarray analysis and were not specifically developed for MS. The most commonly used method is Gaussian random sampling, in which imputations are drawn from a Gaussian distribution centered about the low end of observed quantifications.^20^ This method is employed by the popular Perseus tool^21^ for MS data analysis. In spite of its popularity, this method works poorly.^20^

DreamAI is the best performing current method for TMT imputation.^22^ DreamAI is an ensemble of the six winning methods from the NCI-CPTAC DREAM challenge (https://www.synapse.org/Synapse:syn8228304/wiki/413428), in which more than 20 teams competed to develop imputation methods for CPTAC TMT and isobaric tag for absolute and relative quantification (iTRAQ) data. Being an ensemble method, DreamAI includes high-performing methods such as MissForest.^23^ The DreamAI ensembling strategy outperforms any one of the six individual methods in its ensemble.^22^

Deep learning (DL) has revolutionized our ability to analyze biological data. Most famously, DL is being used to predict protein structures and to discover novel drug targets.^24^ Within the field of proteomics, DL has been used for spectral library generation,^25^ retention time prediction^26^ and peptide de novo sequencing.^27^ One general feature of most DL methods is that they benefit from training with as much data as possible. For example, DL-based de novo sequencing methods have been trained on 30 million peptide-spectrum matches from MassIVE-KB.^27^

In spite of its impressive performance in other domains, DL has not yet gained widespread adoption for proteomics imputation. To our knowledge, there exists only one DL-based proteomics imputation method, called PIMMS, developed for label-free quantification (LFQ).^28^ Perhaps one explanation for the lack of adoption of DL is that existing strategies, including PIMMS, consider only a single dataset at a time and therefore do not benefit from very large training sets derived from multiple MS experiments. In this context underfitting is always a concern, especially when attempting to fit large DL models with many parameters.

Here we present Lupine, a DL-based proteomics imputation method that learns patterns of missingness from multiple datasets simultaneously. We trained Lupine on a joint quantifications matrix that consisted of proteins and MS samples from ten different TMT datasets. Lupine learns low-rank protein and sample embeddings, which are fed into a deep neural network (DNN) to generate predictions. Lupine incorporates an MNAR assumption into its training procedure, explicitly assuming that most missingness is left censored. Our experiments demonstrate that Lupine’s performance improves when trained on ten datasets as opposed to one. Lupine outperforms the current state of the art (DreamAI) in terms of test set accuracy and identifies more differentially expressed (DE) proteins than competing methods. Additionally, Lupine corrects for batch effects in TMT data and learns a meaningful latent representation of proteins and MS samples.

## 2 Results

### 2.1 Overview of the Lupine model

The Lupine model takes as input a partially completed protein-by-sample matrix of quantification values and trains a DNN to complete the matrix by filling in missing values. Lupine has two primary components. First, the model uses two linear embedding layers to learn low-dimensional representations of proteins and MS samples, referred to as protein and sample factors, respectively (Figure 1A). Second, for each missing value, the corresponding protein and sample factors are concatenated and fed into a multilayer perceptron, which generates a protein intensity prediction. The mean squared error (MSE) between model predictions and training set observations is calculated, and the model uses backpropagation to update weights and biases. This process repeats until the model converges. Given the observation that ensembles of individual models, trained with different hyperparameters, often outperform single models,^29^ Lupine is an ensemble of a user-specified number of individual models (default: 10).

**Figure 1.**
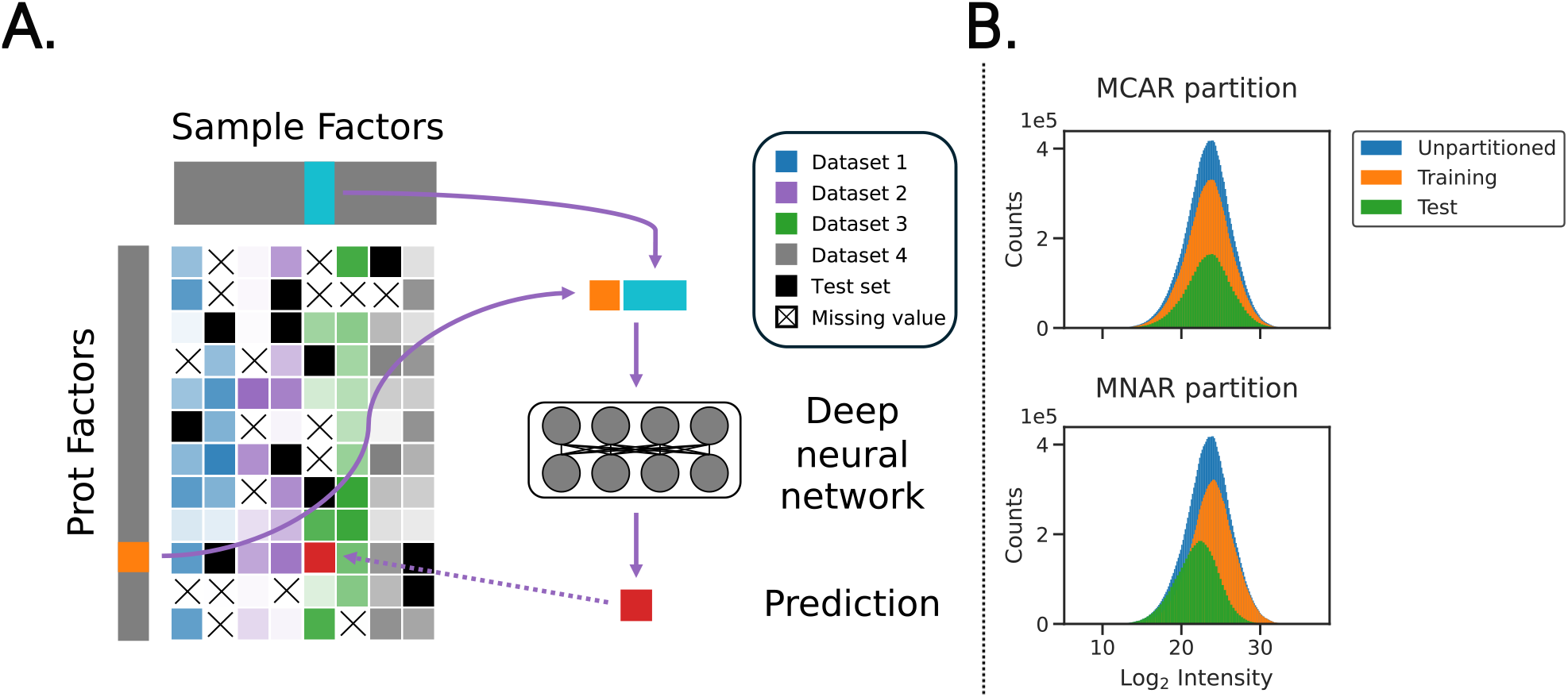
Lupine schematic and data partitioning procedure. **A)** Model schematic. CPTAC datasets were combined into a single joint quantifications matrix to which Lupine was fit. Here the intensity of each color in the joint quantifications matrix corresponds to protein intensity. **B)** Illustration of standard MCAR versus Lupine’s MNAR data partitioning schemes. Lupine was trained entirely in the MNAR setting.

### 2.2 Lupine outperforms the current state of the art for TMT imputation

We hypothesized that, when trained on a large protein-by-samples matrix dervied from many different experiments, Lupine would outperform state-of-the-art methods for MS imputation. Accordingly, we constructed a “joint” quantifications matrix from ten CPTAC datasets (i.e., cohorts) corresponding to ten types of cancer. Rows in the matrix were proteins and columns were de-multiplexed TMT samples. We partitioned this matrix into train and test sets with an MNAR procedure described in Harris et al.^20^ and Section 4.2. This resulted in a test set that was left-skewed relative to the training set (Figure 1B). During model training, we used a “biased” batch selection procedure that preferentially selected matrix entries from the low end of the distribution of intensities for the training set (Supplementary Figure 1). We reasoned that the model would train more effectively if the training set better resembled the test set.

For comparison, we benchmarked Lupine against DreamAI and Gaussian random sampling imputation. DreamAI is the current state of the art for TMT imputation^22^ while Gaussian random sampling is the most commonly used method.^20^ MNAR missingness was simulated and missing values were predicted with each method.

This benchmarking experiment suggests that Lupine indeed outperforms DreamAI and Gaussian random sampling. For all 10 CPTAC cohorts, the test set reconstruction MSE was lower for Lupine than for either competing method (Figure 2A). The residuals between imputed and observed test set values are shown for Lupine and DreamAI in Figure 2B, for three representative cohorts. Points on the diagonal indicate identical predictions made by the two models. Points off the diagonal and centered about 0.0 on the y axis indicate proteins that were more accurately imputed by Lupine. We calculated the fraction of proteins that are more accurately predicted by Lupine than DreamAI. These fractions were 0.641 for CCRCC, 0.614 for HNSCC and 0.606 for LUAD. So as not to bias this analysis by proteins with very low prediction errors, this analysis was limited to proteins for which either method’s residual was *>*0.25.

**Figure 2.**
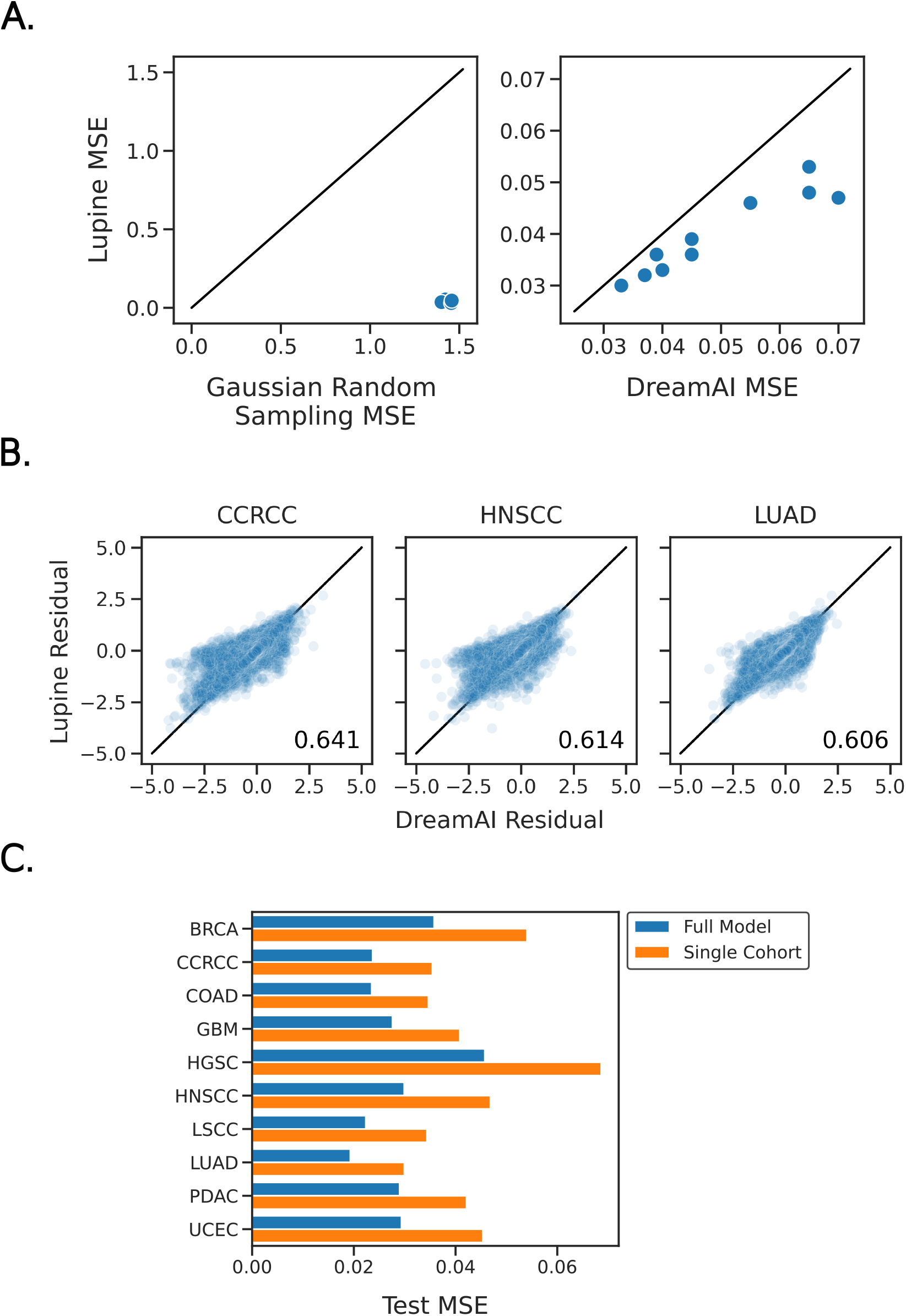
Benchmarking Lupine against state-of-the-art and commonly used proteomics imputation methods. **A)** Test set MSE for Lupine versus Gaussian random sampling (left) and DreamAI (right) imputation, for ten CPTAC cohorts. **B)** Scatterplots of the residuals between model predictions and ground truth (i.e., test set) protein quantifications for Lupine and DreamAI, for three cohorts. The fraction of proteins for which Lupine predictions are more accurate than DreamAI are indicated. **C)** Test set MSE for Lupine models trained on the full joint quantifications matrix versus individual CPTAC cohorts.

Because individual Lupine models are relatively large, containing 4–11M parameters, we hypothesized that Lupine would perform best when applied to larger datasets. We therefore compared the model’s performance when applied to each single cohort versus Lupine applied to the full collection of ten cohorts, using a fixed test set for each comparison. As expected, we observe that Lupine performs better when trained on ten cohorts than on a single cohort (Figure 2C, p=0.0035, paired t-test). On average, the MSE decreases by 34%. This result indicates that including additional MS samples in the training set improves performance even on the original test set samples. This result likely reflects the DL principle that more training data is generally better^29^—the full model may have less of a tendency to overfit.

We performed an ablation experiment to assess the effects of Lupine model ensembling. Models were fit with different random seeds and different numbers of protein factors, sample factors, hidden layers and nodes per hidden layer. We observe a 40% performance gain from ensembling 10 models compared to a single model (Supplementary Figure 2). The performance gain from ensembling 40 models relative to 10 is only 6%. This finding informed our decision to set the default number of models to 10 in the Lupine python package.

It is worth bearing in mind that DreamAI is also an ensembling strategy that averages predictions from six individual imputation methods. Additionally, DreamAI uses a bootstrap aggregation (i.e., bagging) strategy that repeatedly samples the training data, fits models to each bootstrapped set and averages across sets. Thus, it is not unreasonable to compare an ensemble of Lupine models to DreamAI. We attempted to fit DreamAI to the full joint quantifications matrix to include as a baseline in Supplementary Figure 2. However, this proved computationally intractable, because fitting DreamAI to this very large matrix required *>*5 days of compute time (specifically, the MissForest step is extremely time consuming).

### 2.3 Lupine identifies additional differentially expressed proteins

The final goal of quantitative MS experiments is often the identification of proteins that are DE between experimental groups. This step is often hindered by excessive missingness, which can make it especially challenging to identify low-abundance DE proteins and can compromise statistical power, leading to identification of fewer than expected DE proteins. Imputation can help alleviate this problem.

To illustrate Lupine’s practical benefit, we compared the DE proteins identified between tumor and nontumor samples in matrices that had been imputed with several different methods. Lupine was compared to DreamAI, Gaussian random sampling, and no imputation. For each imputed protein, paired t-tests were conducted between tumor and non-tumor samples; proteins with Benjamini-Hochberg adjusted p-values <0.01 and log_2_ fold changes *>*0.5 were considered DE. For seven of eight cohorts, Lupine identifies the most up-regulated proteins, and for six of eight cohorts Lupine identifies the most down-regulated proteins (Figure 3A). Two cohorts, BRCA and GBM, were excluded due to quality control issues with the matched non-tumor samples. These cohorts were also excluded from DE analysis in the original CPTAC publications.^30, 10^

**Figure 3.**
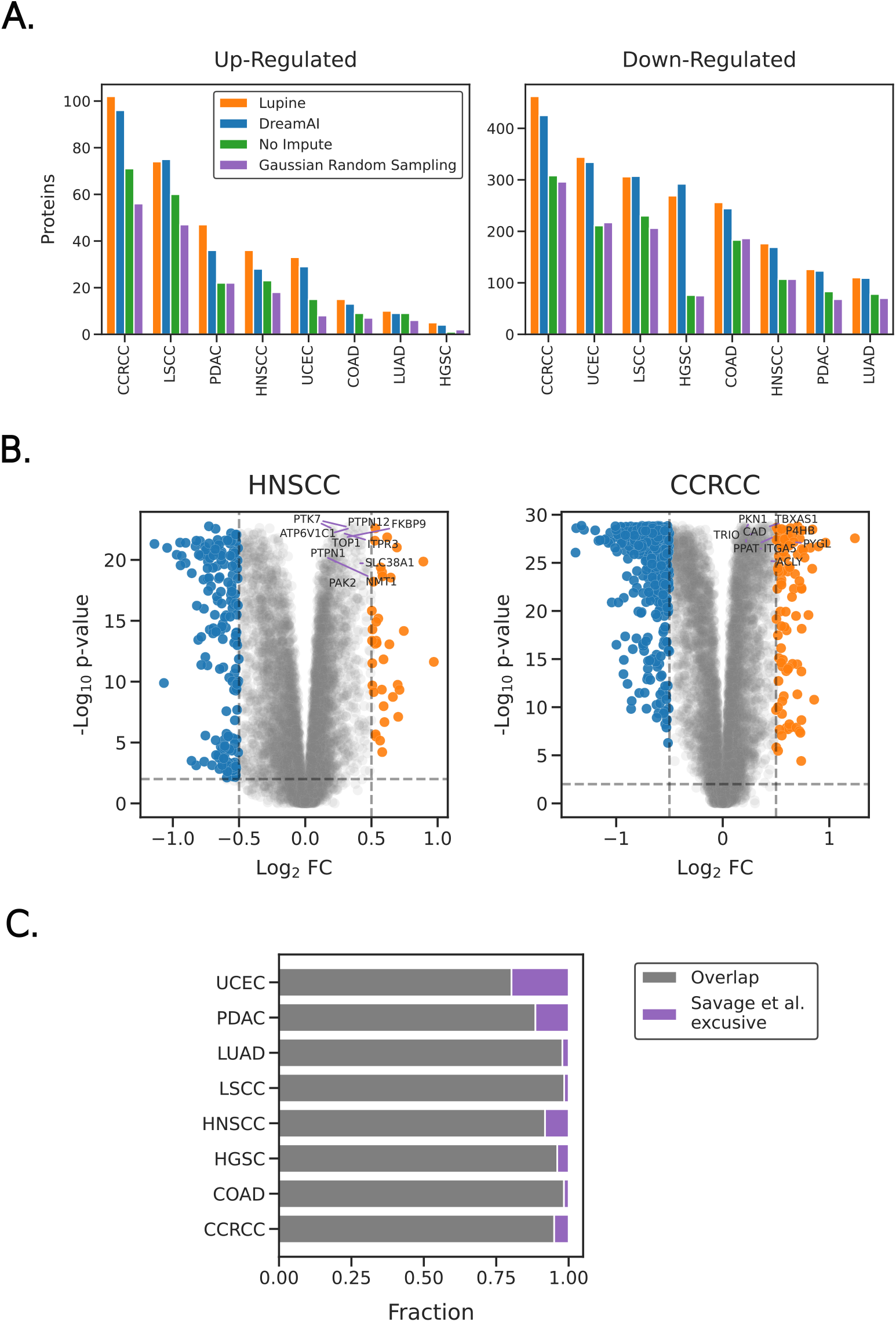
Investigating differentially expressed proteins between tumor and non-tumor samples after imputation. **A)** The number of up- and down-regulated proteins identified within each CPTAC cohort after imputation with Lupine, DreamAI, Gaussian random sampling or no imputation. **B)** Volcano plots of DE proteins imputed with Lupine, for HNSCC and LUAD. Plots have been annotated with the 10 most DE proteins identified by Savage et al.^10^ **C)** The fraction of overlap between DE proteins identified by Savage et al. and those identified after imputation with Lupine.

The DE proteins identified after imputation with Lupine show good concordance with a recent CPTAC publication (Figure 3C). Savage et al. used TMT proteomics data from CPTAC, with no imputation, to identify potential drug targets.^10^ Figure 3C plots the percentage overlap between DE proteins annotated by Savage et al. and our study. For six of eight cohorts the overlap is greater than 90%. This experiment serves as a sanity check, assuring us that expected DE proteins are still identified after imputation with Lupine. Note that the DE protein sets reported by Savage et al. fulfill two criteria: DE between tumor and non-tumor samples and critical for tumor survival and proliferation as determined by a CRISPR knockout screen in cancer cell lines. Figure 3C does not include the set of DE proteins identified by only Lupine because of the possibility that these proteins were also identified as DE by Savage et al. but were not determined essential by their CRISPR screen.

The top ten most DE proteins identified by Savage et al. are also highly DE in our analysis (Figure 3B). Note that Savage et al. did not use log_2_ fold change cutoffs when determining differential expression, which is why some of these top-10 proteins do not show up as DE in our analysis (proteins to the left of the dashed gray lines in Figure 3B). Nevertheless, the p-values from both analyses are among the most significant.

The enriched GO terms for proteins up-regulated in tumor samples are generally related to cell growth and differentiation, DNA replication, and immune system regulation (Table 2). Additional terms include inflammatory response, sugar synthesis, DNA repair, telomere lengthening and cell adhesion.

**Table 1.**
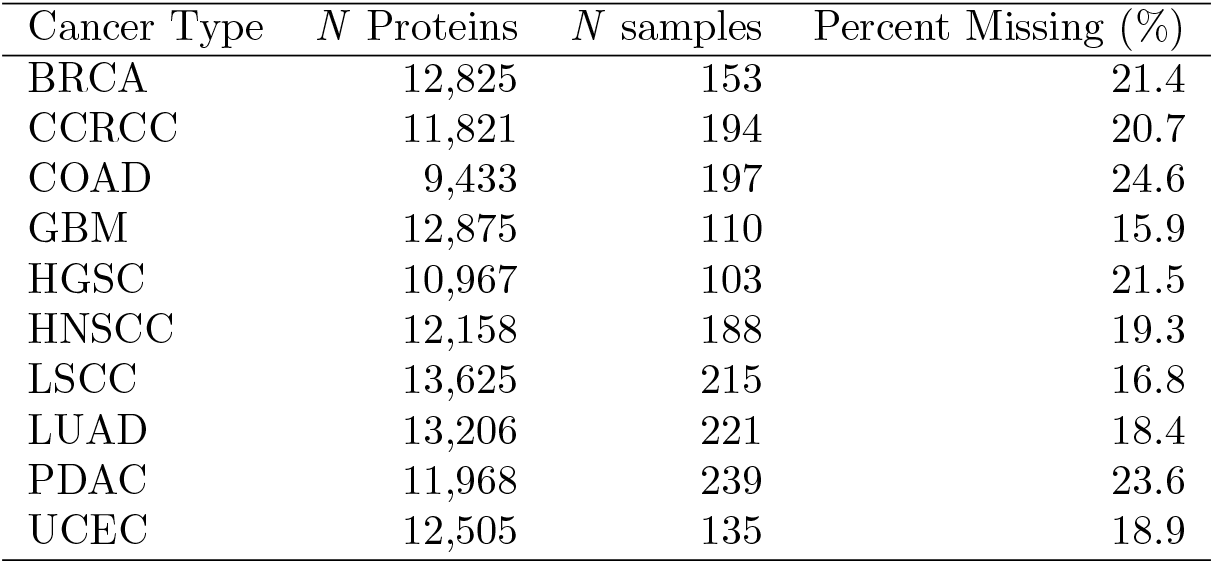
Description of the datasets used in this study. All datasets were generated by the CPTAC Pan-Cancer Proteome project and processed with the pipeline described in the STAR Methods of Li et al.^8^ Abbreviations: BRCA: breast cancer, CCRCC: clear cell renal cell carcinoma, COAD: colon adenocarcinoma, GBM: glioblastoma, HGSC: high-grade serous carcinoma, HNSCC: head and neck squamous cell carcinoma, LSCC: lung squamous cell carcinoma, LUAD: lung adenocarcinoma, PDAC: pancreatic ductal adenocarcinoma, UCEC: uterine corpus endometrial carcinoma.

**Table 2.** Enriched gene ontology (GO) terms among up-regulated proteins for comparisons of tumor versus non-tumor samples after imputation with Lupine.

| Cancer Type | Enriched Gene Sets |
| --- | --- |
| BRCA | Kinetochore assembly, Mitotic spindle assembly checkpoint signaling, Negative regulation of telomere maintenance via telomere lengthening |
| CCRCC | Positive regulation of antigen processing and presentation of peptide or polysaccharide antigen via MHC class II, Negative regulation of T cell co-stimulation, Regulation of tumor necrosis factor (ligand) superfamily member 11 production |
| COAD | Mitotic spindle organization, Regulation of mitotic cell cycle phase transition, Positive regulation of chronic inflammatory response |
| GBM | Sucrose biosynthetic process, Cell-cell adhesion in response to extracellular stimulus, Neutrophil aggregation |
| HGSC | Negative regulation of secretion of lysosomal enzymes, Negative regulation of myeloid progenitor cell differentiation, Mitochondrion migration along actin filament |
| HNSCC | Positive regulation of acute inflammatory response, Neutrophil activation, Regulation of transforming growth factor beta production |
| LSCC | Double-strand break repair via break-induced replication, Mitotic DNA replication initiation, Positive regulation of chromosome condensation |
| LUAD | Cell cycle process, Regulation of transcription by RNA polymerase II, Basophil chemotaxis |
| PDAC | Negative regulation of fibroblast growth factor receptor signaling pathway, Collagen fibril organization, Cell adhesion |
| UCEC | Polyprenol biosynthetic process, Epithelial cell differentiation, Negative regulation of tolerance induction |

Lupine identifies some up- and down-regulated proteins that are not identified by other imputation methods and that may be of biological interest. Lupine identifies on average four additional up-regulated and seven down-regulated proteins that are not identified by DreamAI or Gaussian random sampling imputation (Supplementary Table 1). Most of these proteins are relatively low-intensity (Supplementary Figure 3). These include the protease inhibitor SERPINB7 for LUAD, angiogenesis-associated growth factor VEGFC for PDAC, key developmental transcription factor SOX4 for LSCC, homeobox protein HOXB4 for UCEC, cellcell adhesion glycoprotein CDH4 for CCRCC and Ras oncogene family member RAB40C in HNSCC.

However, we stress that the additional DE proteins identified by Lupine should be considered in the context of hypothesis generation. For experiments that are tolerant of false positives—for example, screening small molecules for therapeutic potential—Lupine’s ability to identify DE proteins may prove invaluable. But the gold standard for differential expression remains the directly observed quantifications from lower-throughput targeted assays.

### 2.4 Within-complex correlations are higher than non-complex correlations

Many proteins function within larger protein complexes and should thus exhibit correlated abundances. In principle, a poor imputation method could introduce noise and thus degrade these correlations. Accordingly, we tested whether within-complex correlations are higher than non-complex correlations both before and after imputation with Lupine. We used protein complex annotations from the CORUM^31^ database and limited this analysis to proteins with initial missingness fractions <50% in the joint quantifications matrix.

For each cohort, we calculated the Spearman correlations of intensities for all pairs of proteins within the same CORUM complex, and for the same number of pairs of randomly selected proteins.

Within-complex correlations are significantly higher than non-complex correlations both before and after Lupine imputation (Figure 4, paired t-test p-values <0.01). For Lupine imputed data, the mean within-complex correlation is 0.270, compared to 0.003 for non-complex. For unimputed data, the within-complex correlation is 0.284, compared to 0.001 for non-complex. It is reassuring that Lupine does not meaningfully reduce within-complex correlations and that non-complex correlations remain ∼ 0.0 after imputation. This result suggests that Lupine is not learning spurious correlations between proteins.

**Figure 4.**
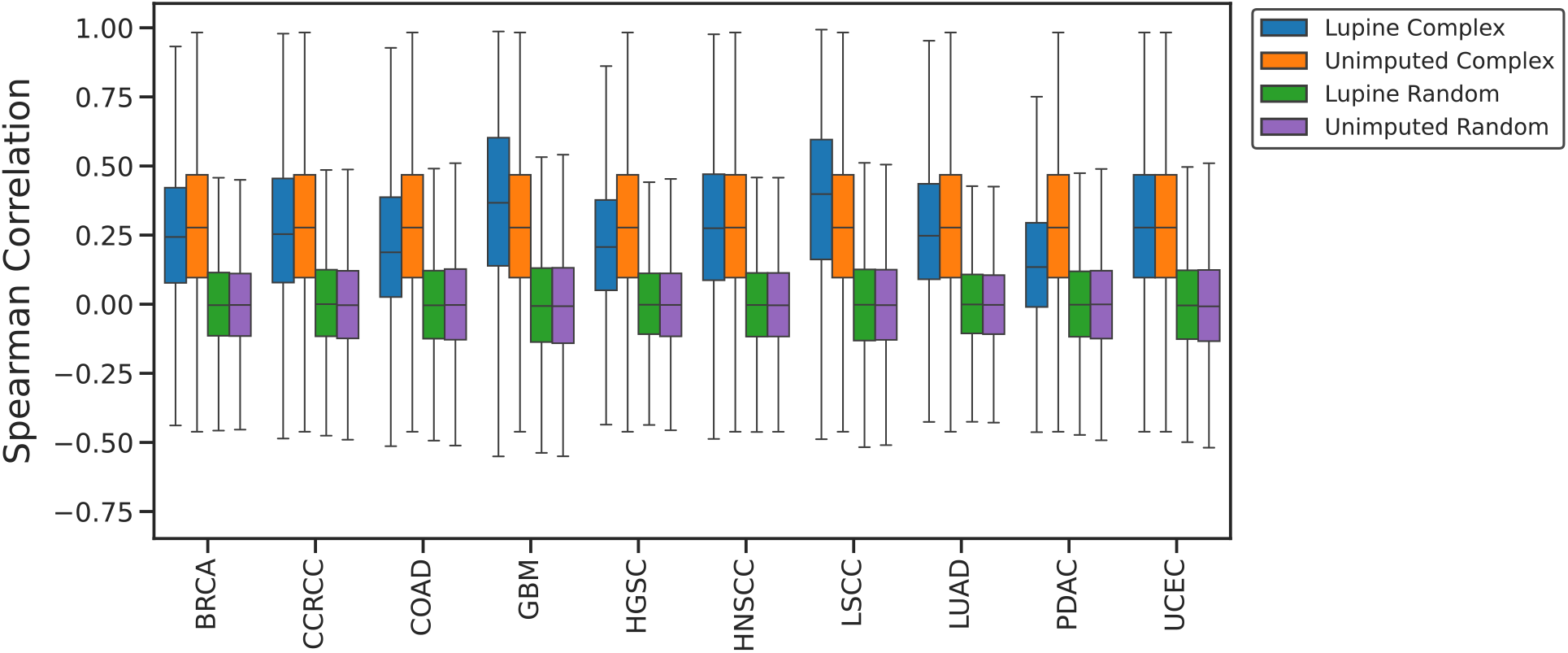
Within-complex versus non-complex protein-protein correlations before and after imputation. Distributions of Spearman correlations of intensities of pairs of proteins potentially in the same complex (blue and orange) versus pairs of randomly selected proteins (green and purple), before and after imputation with Lupine.

### 2.5 Lupine corrects for batch effects in TMT data

Batch effects represent a major challenge to the interpretation of TMT data.^6^ Batch effect correction is often performed with methods like Combat^32^ and surrogate variable analysis (SVA).^33^ While Lupine is not a batch correction method and is not intended to replace Combat or SVA, we have found that Lupine appears to automatically carry out some batch correction, even without having access to any batch structure annotations.

For a single cohort, CCRCC, we imputed missing values with Lupine and column minimum (herein “min”) impute, another naive but commonly used method.^20^ Column min consists of replacing missing values with the lowest observed quantification for a given MS sample. We then performed dimensionality reduction with UMAP (https://umap-learn.readthedocs.io/en/latest/) and colored by sample type annotations and TMT batch IDs (Figure 5). A priori, we did not expect column min to correct batch effects—we would have preferred to perform this analysis on unimputed quantifications, but UMAP requires complete matrices so we had to apply some imputation procedure.

**Figure 5.**
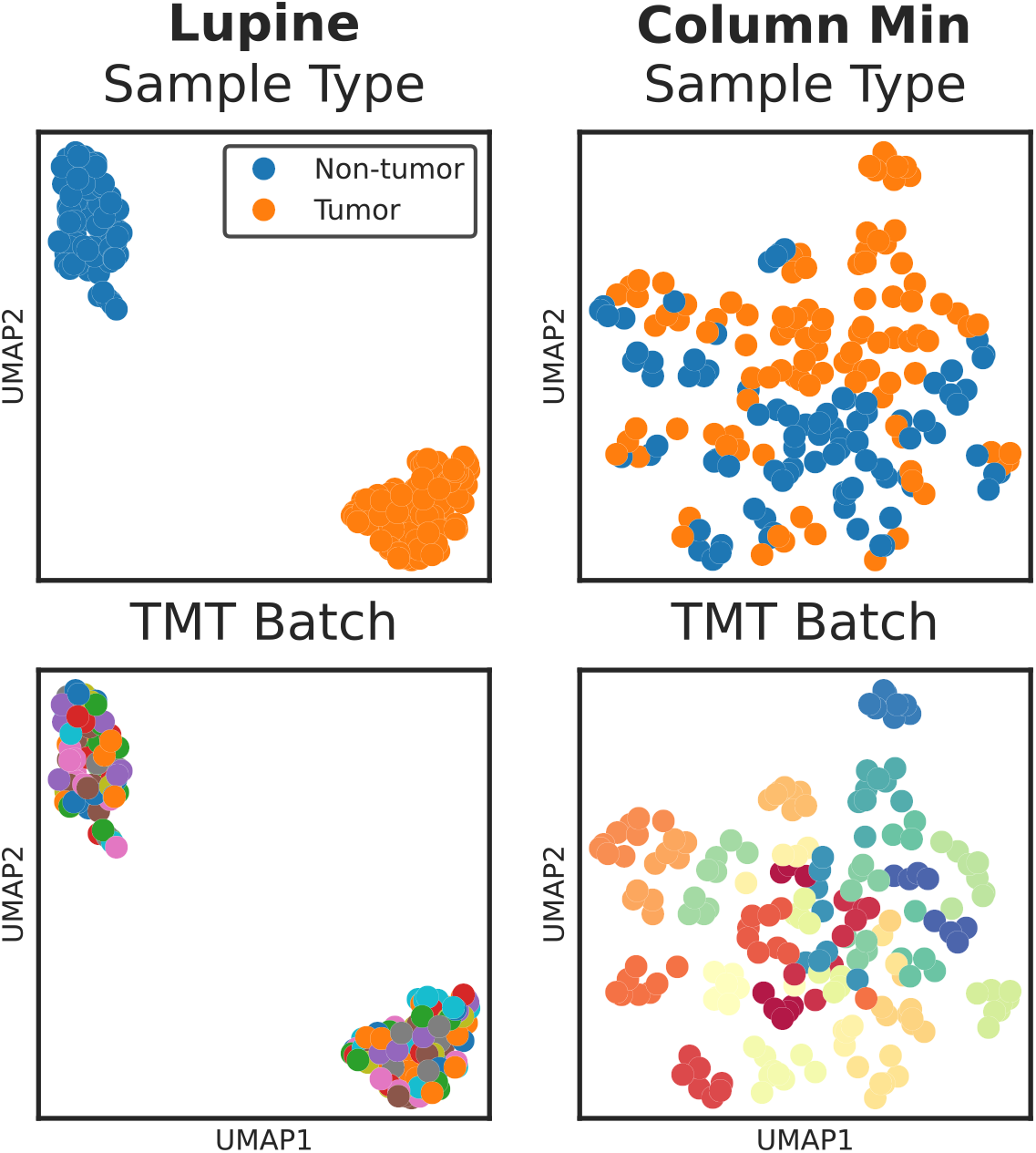
UMAP projections of protein intensities for a single CPTAC cohort following imputation with Lupine or column min. Top panel is colored by sample type annotation (i.e., tumor versus non-tumor status); bottom panel is colored by TMT batch ID.

Intriguingly, we observe that the Lupine imputed proteins cluster by sample type, whereas the column min imputed proteins cluster primarily by TMT batch (Figure 5). The UMAP projection shown in the left panel of Figure 5 closely resembles the PCA analysis conducted by Clark et al. in CPTAC’s original description of the CCRCC data.^34^ Importantly, Clark et al. applied ComBat in addition to imputation, whereas we simply imputed with Lupine. The majority of the cohorts cluster by sample type following Lupine imputation (Supplementary Figure 4). Additionally, we see remarkable separation of CPTAC cohorts and tumor versus non-tumor samples when looking at the entire joint quantifications matrix following Lupine imputation (Supplementary Figure 5). Thus, Lupine apparently learns to separate MS samples according to biological rather than technical signal.

We do not suggest that Lupine can serve as a replacement for batch effect correction algorithms like ComBat and SVA. For one thing, these methods differ from Lupine in that they are supervised—the identities of the batches are provided as input to the method. Lupine learns to separate technical from biological signal purely from the data themselves. For this reason we do not compare Lupine to ComBat or SVA. But the result in Figure 5 is interesting nonetheless and will be the subject of future exploration. Future versions of Lupine may be geared toward imputation and batch correction for single cell proteomics, a domain in which batch effects and missing values currently represent significant barriers to analysis.^35, 36^

### 2.6 Lupine learns a representation of proteins and MS samples

Lupine, like many DL models, learns in part by projecting observed inputs into a latent embedding space. We hypothesized that, if Lupine is learning effectively, then this embedding space should correspond to some aspect of biological signal rather than noise. We examined the latent embeddings of a Lupine model fit to the joint quantifications matrix by reducing the embedding dimensionality with UMAP and then coloring the 2D projection using various types of metadata. We found that Lupine’s sample embeddings cluster exclusively by CPTAC cohort, with separate clusters for tumor and non-tumor samples (Figure 6). On the other hand, the protein embeddings do not form separate clusters but exhibit a clear gradation according to protein missingness fraction. Importantly, Lupine does not have access to any metadata; instead, these relationships are learned directly from the protein quantifications. We also investigated coloring the protein embeddings by various physicochemical properties such as size, charge, hydrophobicity, etc., but this analysis did not reveal any clear trends.

**Figure 6.**
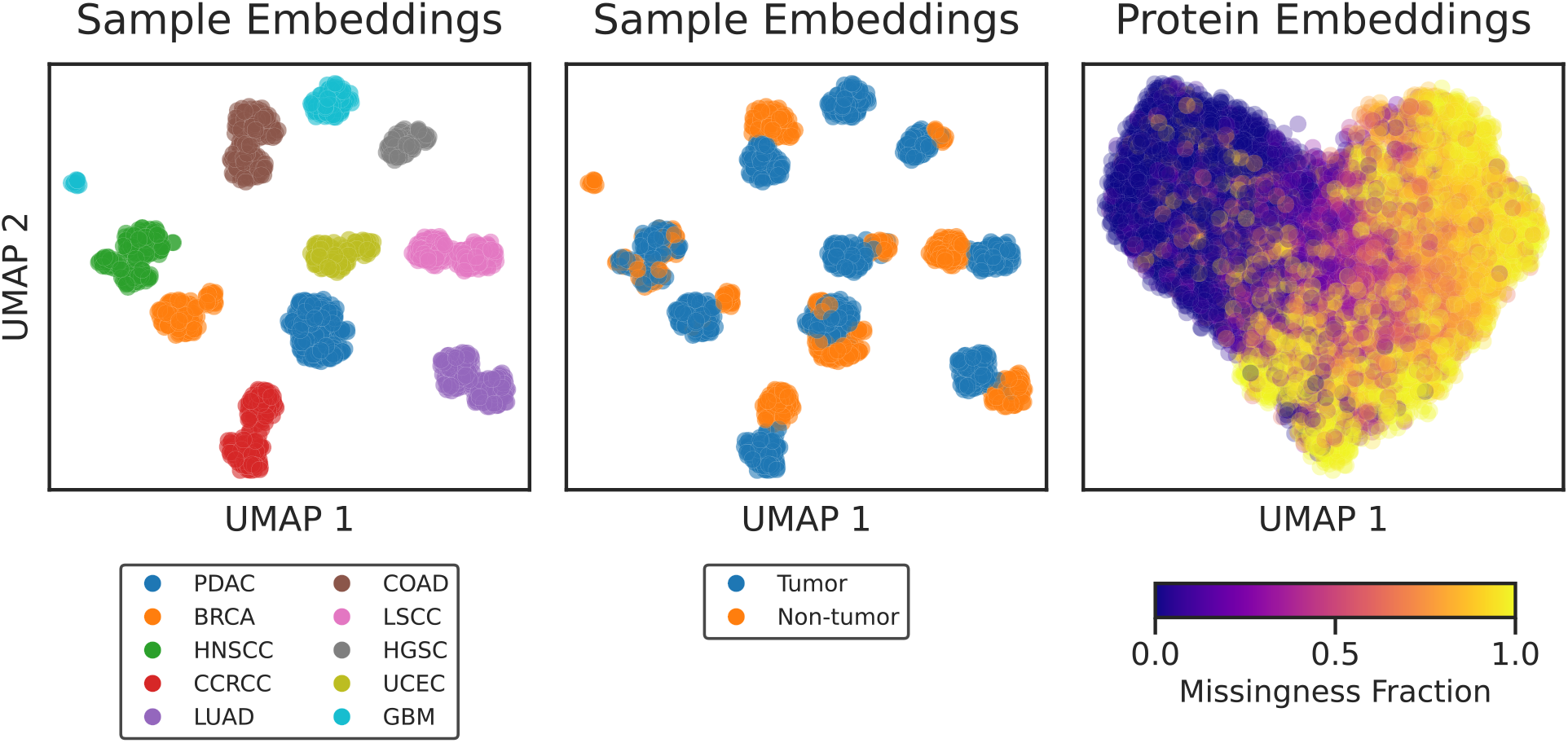
UMAP projections of Lupine’s sample and protein embeddings. Left: points correspond to patient samples and are colored by CPTAC cohort. Center: points correspond to patient samples and are colored by sample type. Right: points correspond to proteins and are colored by protein missingness fraction in the joint quantifications matrix.

Future work will continue to explore this learned embedding space. Several interesting questions may be addressed, including: Are there outlier patient samples that seem to cluster disconcordantly from their clinical annotations? What can we learn about the biology of such outliers? What are the additional properties that drive Lupine’s protein embeddings and what can they tell us about protein missingness?

## 3 Discussion

Lupine is a proteomics imputation method that learns from multiple datasets at once. Lupine was trained on a joint protein quantifications matrix consisting of ten separate datasets for a total of 18,162 proteins across 1,755 MS samples. Our experiments demonstrate that Lupine performs better when trained on this joint quantifications matrix than on any of the ten individual datasets (Figure 2C). We speculate that a Lupine model trained on an even larger training set will perform even better. Lupine is distinct among imputation methods in that it is designed to learn from many experiments; existing methods consider a single MS experiment at a time.

It is worth noting, however, that here Lupine was trained on a rather homogenous joint quantifications matrix. All MS samples were derived from cancer patients, were run on Thermo Fisher instruments, and were processed with a common data analysis pipeline. In the future, Lupine will be fit to a more heterogenous training matrix that includes LFQ and data-independent acquisition (DIA) MS samples, as well as non-cancer samples. We speculate that given the model’s batch effect correction capacity (Figure 5), Lupine will learn the differences between MS acquisition strategies and still emphasize biological signal.

Lupine is a Python package available on GitHub (https://github.com/Noble-Lab/lupine) with an MIT open source license. The Lupine package includes a command that allows users to attach their own TMT MS samples to the existing joint quantifications matrix. Lupine may then be fit to the modified joint quantifications matrix and will impute the user’s TMT samples. The package’s documentation provides guidelines for selecting model hyperparameters. Currently, model training requires a GPU, but we are working on a distilled model that can be run on CPU in a reasonable timeframe. Given the relationship between performance and the number of ensembled models (Supplementary Figure 2), the Lupine package includes a parameter specifying the number of models to ensemble, with a default of 10. Additionally, the Lupine imputed CPTAC protein quantification matrices produced by this study are available on Zenodo (https://zenodo.org/records/13146445). We hope that the community continues to mine these imputed MS samples for insights into cancer biology.

Future versions of Lupine will incorporate patient-matched multi-omic measurements from CPTAC including phosphoproteomics, whole exome sequencing, transcriptomics and DNA methylation. Lupine’s imputation framework will be extended to encompass these other data modalities, allowing the model to learn a rich embedding space that captures DNA, RNA and protein measurements. Lupine’s embedding space will then be mined for insights into cancer biology.

Additionally, future versions of Lupine will be trained on peptide rather than protein quantifications. We initially trained Lupine on protein quantifications because the majority of MS experiments report protein-level identifications and quantifications. However, protein roll-up can mask important biological signal, for example, in the context of neurodegenerative diseases that are driven by aberrant proteoforms.^37, 38^ Accordingly, future versions of Lupine will focus specifically on peptide imputation.

Finally, future work will focus on attaching statistical measures of confidence to imputed values. Because imputed values were not directly measured by the instrument, they should be thought of as lower quality measurements than observed values. However, most researchers do not take this into account when performing downstream analysis, and instead treat imputed and observed values the same. Prediction-powered inference (PPI) is a statistical framework for attaching confidence intervals to DL or machine learning predictions.^39^ A powerful extension of this work is to build a PPI framework for Lupine imputation that allows researchers to prioritize higher confidence observed values in downstream analysis, while still benefiting from the increased statistical power and low-abundance quantifications afforded by imputation.

## 4 Methods

### 4.1 Joint quantifications matrix construction

We obtained TMT proteomics data—collected by CPTAC—through the Proteomics Data Commons web portal (https://pdc.cancer.gov/pdc/cptac-pancancer, file name: Proteome UMich GENCODE34 v1.zip). Data were collected at five centers: Pacific Northwest National Laboratories, Vanderbilt University, Johns Hopkins University, Harvard Medical School and the Broad Institute. Samples were run on Thermo Fisher orbitrap instruments. Datasets were processed with the same analytical pipeline, the full details of which are provided in the STAR Methods of Li et al.^8^ Briefly, this pipeline consisted of peptide search with MSFragger^40^ against a GENCODEv34 database and post-processing with Philosopher^41^ and TMT-Integrator.^40^ TMT-10 or -11 plexes were normalized to a common reference channel then summarized at the protein level.

We constructed a joint quantifications matrix from CPTAC datasets. Rows in the joint quantifications matrix were proteins and columns were de-multiplexed TMT samples. When adding new MS samples (i.e., columns) to the matrix, if a protein was not quantified in the given sample, then the matrix entry was assigned a value of NaN (not a number) to indicate a missing value. We removed proteins quantified in fewer than 18 samples. Sample IDs containing the following key words were excluded from analysis: “RefInt”, “QC”, “pool”, “pooled”, “reference”, “NCI”, “NX”, “ref”. After filtering, this dataset consisted of 18,162 proteins and 1,755 samples. The overall missingness fraction was 48.7%.

### 4.2 Dataset partitioning and batch selection

We used an MNAR data partitioning procedure to simulate the type of missingness most common to MS proteomics.^1, 2, 12^ Given our joint quantifications matrix *X* with rows *i* and columns *j*, we constructed a “thresholds” matrix *T*_*i*×*j*_. *T* was populated with values sampled from a Gaussian distribution centered about the 25^th^ percentile of *X* and with standard deviation (*σ*): 1.1 × *X*_*σ*_. For each *X*_*ij*_, if the corresponding *T*_*ij*_*>X*_*ij*_, then *X*_*ij*_ was assigned to the training set. Otherwise, a Bernoulli trial with success probability 0.61 was conducted. If the trial was successful, then *X*_*ij*_ was assigned to the test set, otherwise, *X*_*ij*_ was assigned to the training set. The Bernoulli success probability and Gaussian distribution mean and standard deviation were tuned such that 20% of present matrix entries were assigned to the test set and the remaining 80% were assigned to the training set.

We implemented a “biased” batch selection procedure during model training. To achieve this we conducted multiple rounds of the Bernoulli trial-based selection procedure described above, in which the training set was sampled with replacement to create a series of training batches. Low-intensity proteins were preferentially selected for training batches (Supplementary Figure 1). The resulting distribution of training batches is left-skewed relative to the whole training set. This procedure was undertaken without looking at the test set.

### 4.3 Model implementation

Lupine uses a DNN to learn a low-dimensional representation of proteins and MS samples (Figure 1A). The protein and sample factor matrices, referred to as *W* and *H* respectively, were randomly initialized linear embedding layers. The shapes were as follows: *W* : (18,162, *p*) given *p* protein factors and *H*: (*s*, 1,755) given *s* sample factors. *W* and *H* contained no missing values. Selection of model hyperparameters *p* and *s* is described in the next section.

The Lupine training procedure was as follows. For each training batch, for each *Xtrain*_*i,j*_, the corresponding protein (*W*_*i*_) and sample (*H*_*i*_) factors were concatenated and fed into a fully-connected multilayer perceptron (i.e., the DNN). This DNN consisted of a variable number of hidden layers and nodes per layer, with leaky ReLU activations (negative slope 0.1) after each layer. The DNN output a predicted value *Xpred*_*i,j*_. The mean squared error (MSE) between *Xtrain*_*i,j*_ and *Xpred*_*i,j*_ was calculated. This procedure was repeated for every *Xtrain*_*i,j*_ in the train batch, then model parameters *W, H* and the DNN weights and biases were updated with backpropagation. The Adam optimizer was used, with a learning rate of 0.001. The training batch size was 128.

A validation set was used to determine model convergence. This validation set consisted of 10% of matrix entries randomly selected (MCAR) from the training set. Training proceeded until one of two stopping criteria were triggered, as evaluated on the validation set: (1) The ratio of the difference between the best loss and the current loss (numerator) to the best loss (denominator) was calculated at the end of each epoch. If this ratio was less than a tolerance parameter (0.001) for 10 successive epochs, then training stopped. (2) A Wilcoxon rank sum test was conducted between two “windows” of validation set MSEs, window 1 being the previous five training epochs and window 2 being training epochs *n* − 10 to *n* −15, where *n* is the current epoch. If the one-sided Wilcoxon p-value comparing window 2 to window 1 was <0.05, then training stopped, indicating that validation error had stopped decreasing and had started to increase.

Lupine was implemented in PyTorch v1.10.2 and was trained on a combination of nVidia GeForce GTX 1080, GeForce RTX 2080 and GeForce RTX 2080 Ti graphic processing units (GPUs).

## 4.4 Benchmarking

The joint quantifications matrix was partitioned into 80% train and 20% test with an MNAR procedure, as described in Section 4.2. Proteins with fewer than 18 present values in the training set were removed from the training and test matrices. Lupine was fit to this training set.

The training set was then divided into 10 subsets, each containing the MS samples for a single CPTAC cohort. For each cohort subset, proteins with fewer than three present values were removed from the matrix. DreamAI and Gaussian random sampling imputation were fit to each cohort subset. DreamAI was accessed via GitHub (https://github.com/WangLab-MSSM/DreamAI) and was run in R. Gaussian random sampling was implemented with custom python code replicating the Perseus procedure described here: https://cox-labs.github.io/coxdocs/replacemissingfromgaussian.html.

Lupine is an ensemble model consisting of *n* individual models trained with different values for the following hyperparameters: number of protein factors *p*, number of sample factors *s*, number of hidden layers and number of nodes per hidden layer. The search space for each hyperparameter was as follows: *p* and *s*: [64, 128, 256, 512, 1024], hidden layers: [1, 2, 4], nodes per hidden layer: [512, 1024, 2048]. Of the 225 possible combinations of hyperparameters, a random selection of 42 was chosen. *n* = 42 independent Lupine models were then fitted to the full training matrix. Each model used a different random seed and partitioned a different 10% of training set *X*_*ij*_s into its validation set. The predictions from these 42 models were then averaged to generate the final Lupine reconstructed matrix.

To enable one-to-one performance comparisons, for each cohort, the Lupine reconstructed matrix was subset to include only the proteins contained by the DreamAI/Gaussian random sampling training matrix for that cohort. In this way we evaluated predictions on the same set of proteins and MS samples for all three models. We calculated the MSE between model predictions and the observed test set values for each cohort.

### 4.5 Differential expression

We identified DE proteins between tumor and non-tumor samples within each CPTAC cohort after imputation with three methods. Lupine, DreamAI and Gaussian random sampling imputation were performed as described in the previous section. CPTAC metadata, containing sample type annotations, were obtained from the CPTAC data portal: https://pdc.cancer.gov/pdc/. For each cohort, proteins with *>*50% missingness prior to imputation were excluded from DE analysis.

For each imputed protein, paired t-tests were conducted between protein quantifications from tumor and non-tumor samples. We also included a no imputation condition, in which missing values were ignored when conducting t-tests. P-values were adjusted with the Benjamini-Hochberg (BH) procedure.^42^ Proteins with BH adjusted p-values <0.01 and log_2_ fold changes *>*0.5 were considered DE. This same statistical procedure was used to identify DE proteins by CPTAC in Savage et al.^10^ The exception is that Savage et al. omitted the log_2_ fold change criteria that we use here.

To identify enriched gene ontology (GO) terms, we used the PANTHER overrepresentation test (https://pantherdb.org/tools/compareToRefList.jsp). PANTHER19.0 was used. For each cohort, the set of up-regulated DE proteins was compared to a background list consisting of all 18,162 proteins in the joint quantifications matrix. The test type was Fisher’s exact and the annotation dataset was “GO biological process complete.” Representative enriched GO biological processes were chosen to populate Table 2. This analysis was conducted after imputation with Lupine.

### 4.6 Protein complex analysis

We compared within-complex to non-complex protein-protein correlations. The first step was to convert ENSEMBL v44 protein IDs to HGNC IDs within our joint quantifications matrix. We then divided our Lupine imputed joint quantifications matrix into 10 subsets according to CPTAC cohort. Proteins with initial (pre-imputation) missingness fractions *>*0.5 were removed from these matrices. Each cohort matrix was then subset to only tumor samples. We then searched HGNC IDs against the CORUM database (https://mips.helmholtz-muenchen.de/corum/#download, file name: humanComplexes.txt) of known human protein complexes^31^ for each cohort. For each CORUM annotated complex, we computed the Spearman correlations between every pair of subunits in the complex. We then computed the same number of Spearman correlations for pairs of randomly selected proteins from the cohort matrix.

## Data Availability

Unimputed CPTAC datasets were obtained from the Proteomics Data Commons web portal (https://pdc.cancer.gov/pdc/cptacpancancer). The name of the file we accessed was Proteome UMich GENCODE34 v1.zip. The Lupine imputed versions of these protein quantifications are available at https://zenodo.org/records/13146445.

## Code Availability

The code used to generate all the figures in this manuscript can be found at https://github.com/Noble-Lab/2023_harris_deep_impute. A python package implementing Lupine can be found at https://github.com/Noble-Lab/lupine.

## Acknowledgments

The authors thank Michael J. MacCoss and Bo Wen from Genome Sciences for project guidance and help with all things related to CPTAC, respectively. We thank Samuel H. Payne from Brigham Young University for pointing us to the relevant datasets and William E. Fondrie from Talus Bioscience for initial project guidance. This work was supported by National Institute on Aging grant number 1F31AG082395-01 and by NSF award 2245300.

## Author contributions

Project was conceived of by W.S.N. Experiments were performed, text was written and figures were generated by L.H.

