## Supplement for "Imputation of cancer proteomics data with a deep model that learns from many datasets"

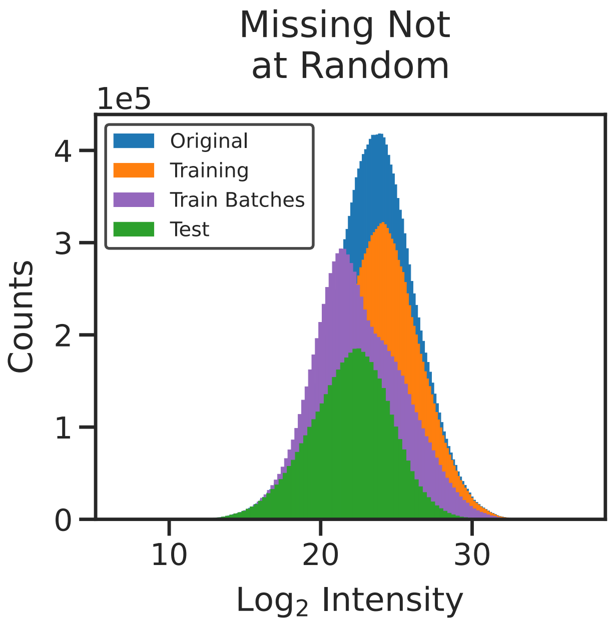


Supplementary Figure 1. **Lupine used a biased batch selection procedure during model training.** The joint quantifications matrix (blue) was partitioned into training (orange) and test (green) sets with an MNAR procedure described in Section 4.2. Training batches were preferentially selected from the low-end of the training set distribution; repeated sampling of the training set was allowed.

| Cohort | Up-regulated | Down-regulated |
|---|---:|---:|
| CCRCC | 6 | 24 |
| COAD | 2 | 7 |
| HGSC | 1 | 7 |
| HNSCC | 7 | 2 |
| LSCC | 2 | 0 |
| LUAD | 1 | 2 |
| PDAC | 7 | 5 |
| UCEC | 5 | 7 |

Supplementary Table 1. **Counts of differentially expressed proteins uniquely identified after imputation with Lupine.** Differential expression testing was performed as described in Section 4.5 after imputation with Lupine, DreamAI or Gaussian random sampling. Numbers of DE proteins uniquely identified after Lupine imputation are indicated.

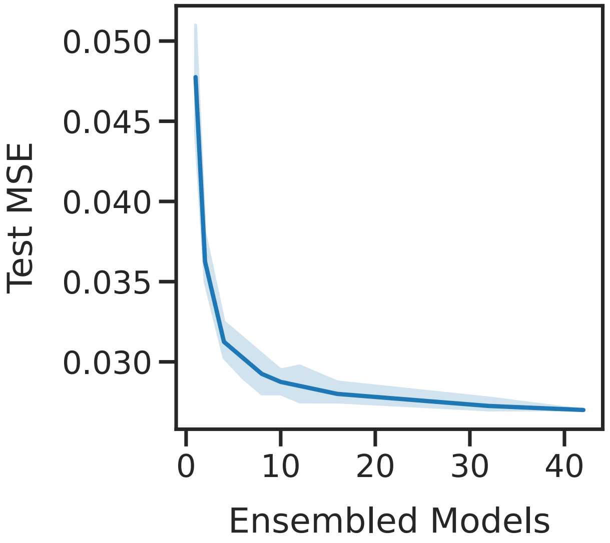


Supplementary Figure 2. **Test set MSE as a function of the number of ensembled Lupine models.** Lupine models were fit to the training set derived from the joint quantifications matrix. Four sets of ensembled models were fit to obtain the 95% confidence intervals shown.

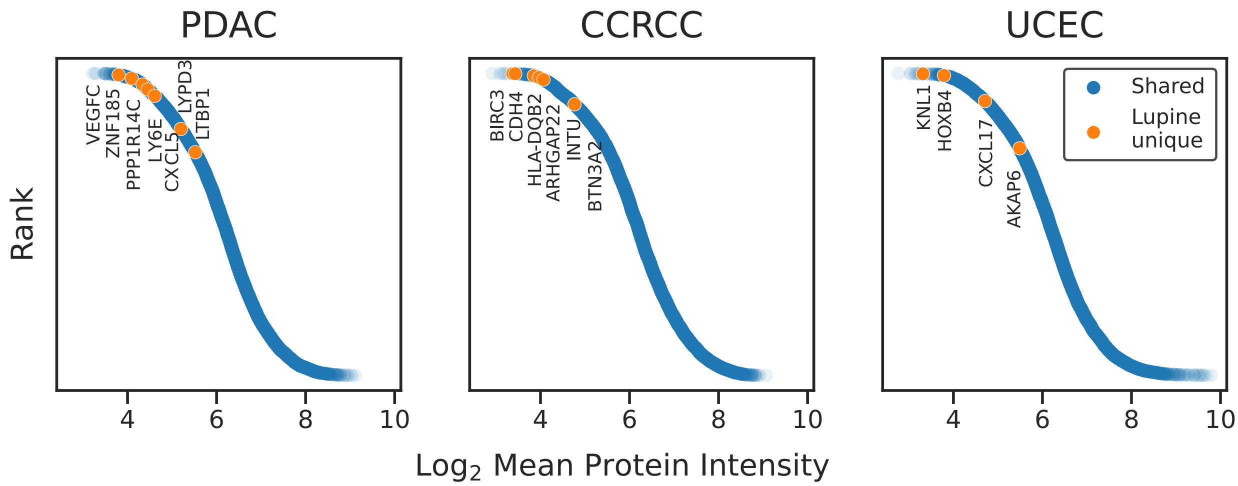


Supplementary Figure 3. **Rank plots of mean intensities of differentially expressed proteins identified between tumor and non-tumor samples.** Blue indicates DE proteins identified after imputation with at least two of Lupine, DreamAI and Gaussian random sampling. Orange indicates DE proteins uniquely identified after Lupine imputation. The mean protein intensities for non-tumor samples are shown. Proteins with >90% initial missingness were excluded from this analysis.

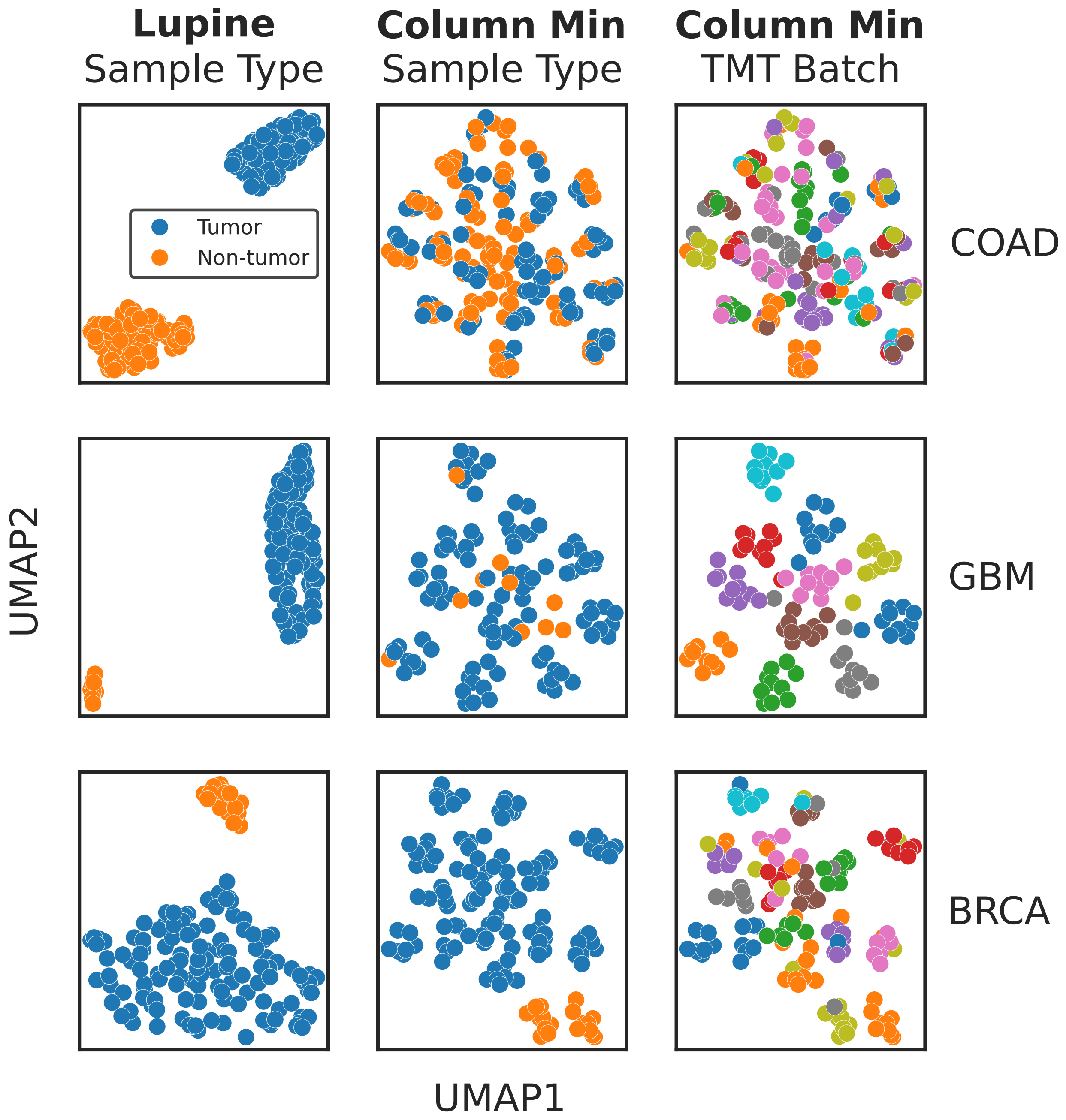


Supplementary Figure 4. **UMAP projections of protein intensities following imputation for three additional CPTAC cohorts.** Left panel: Lupine imputation colored by sample type; center panel: column min imputation colored by sample type; right panel: column min imputation colored by TMT batch ID.

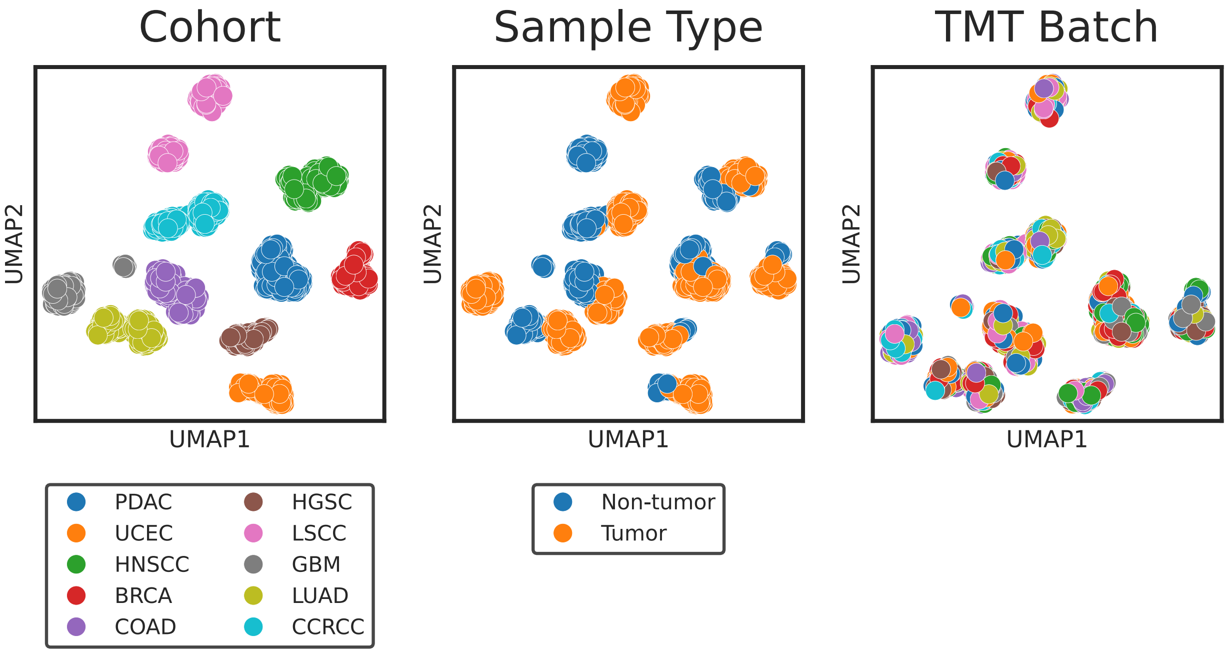


Supplementary Figure 5. **UMAP projections of the Lupine imputed joint quantifications matrix.** Left: colored by CPTAC cohort; center: colored by sample type; right: colored by TMT batch ID. Each point corresponds to an MS sample.
